# Mucosal microbiota and metabolome along the intestinal tracts reveals location specific relationship

**DOI:** 10.1101/454496

**Authors:** Ce Yuan, Melanie Graham, Christopher Staley, Subbaya Subramanian

## Abstract

**Background:** The intestinal microbiota has been recognized as an important component for maintaining human health. The perturbation to its structure has been implicated in many diseases, such as obesity and cancers. The microbiota is highly metabolically active and plays a role in many metabolic pathways absent from the human host. Altered microbiota metabolism has also been linked to obesity, cardiovascular disease, and colorectal cancer. However, there is a gap in the current knowledge of how the microbiota interacts with its host. Here we performed an integrated analysis between the mucosal-associated microbiota and the mucosal tissue metabolomics in healthy non-human primates (NHPs) to investigate these relationships.

**Results:** We found that the overall microbiota composition is influenced by both the tissue location as well as the host individual. The NHPs intestinal microbiota predominantly comprised of members of the phyla Firmicutes, Bacteroidetes, and Proteobacteria. The large intestines contain more Spirochaetes, Tenericutes, and Lentisphaera phyla members. The small intestinal tissues have no significantly different microbiota compositions, while the cecum and distal colon differ greatly in the microbiota compositions. The metabolomics profile reveals a total of 140 metabolites with different concentration between the small and large intestines. The correlations between microbiota and tissue metabolites showed a dense and interconnected network in the small intestines while a sparse network in the large intestines.

**Conclusions:** Our analysis revealed an intricate global relationship between the microbiota and the host tissue metabolome that is mainly driven by the distal colon. Most importantly, we found location specific microbiota-metabolite correlations that have potential implications for studying host-microbiota metabolic interactions.

## Background

The human gastrointestinal (GI) tract harbors trillions of microorganisms, including thousands of bacterial species, termed the microbiota [1]. It has become evident that the gut microbiota is important in regulating and maintaining the health of the host and is implicated in many diseases, such as cancers [1–5]. Despite numerous studies indicating important roles of microbiota in diseases, many of these studies have largely focused on the taxonomic composition of the microbiota [6]. The underlying metabolic features associated with the host-microbiota interaction, however, still remains unclear.

Previous studies suggest the gut microbiota produces a vast amount of metabolites. Some metabolites, such as vitamin B, vitamin K, and short-chain fatty acids (SCFAs), are essential to maintaining homeostasis of the colon epithelial cells [7–9]. Moreover, the metabolic interaction between the host and its microbiota has widespread implications throughout the body [8]. For example, the obesity-associated microbiota has been shown to possess an increased metabolic capability to harvest energy from food [10, 11], and the metabolism of L-carnitine by the gut microbiota has been shown to promote atherosclerosis [12]. These studies suggest potential metabolic shifts of the microbiota either in response to or responsible for the host metabolic state [11].

The most direct and active metabolic interactions between the host and its microbiota occur in the large intestines. Most notably, the vast majority (~70%) of energy required by the normal colon epithelium comes from butyrate produced by the microbiota through fermentation of complex carbohydrates [13]. Without a functional microbiota, the colon epithelia undergo autophagy and fail to maintain normal structure and function [14]. More importantly, these metabolic interactions have important implications in colorectal cancer, the second most deadly cancer in the United States [1, 5, 7, 8, 15–18]. Furthermore, bile acids, produced by the liver and modified by the intestinal bacteria, have been shown play signaling roles that impact the progression of obesity and the metabolic syndrome [19]. Similarly, short-chain fatty acids produced from the microbial breakdown of dietary fiber serve as signaling molecules that can affect host metabolic responses in distant organs [20].

The specific mucosal host-microbiota metabolic interactions occurring along a healthy human GI tract are largely unknown. Although the microbiota and metabolome variations along the GI tract has been investigated in rodents and other animals, the dietary and anatomical differences between humans and these animals render these data less informative for humans [21–26]. Here, we investigated the microbiota and metabolome profiles along the intestinal tracts of healthy baboons (*Papio anubis*), a family of Old World monkeys. We collected tissue samples from the duodenum, jejunum, ileum, cecum, proximal colon and distal colon. 16S rRNA gene sequencing (16S-Seq) was performed to identify the mucosal surface microbiota composition. We also used untargeted metabolomics analyses on the immediately adjacent tissues to profile the tissue metabolites. These data serve to more comprehensively establish the intestinal host-microbiota metabolic interactions in non-human primates.

## Results

### Microbiota diversity and composition along the baboon GI tract

We first assessed the baboon GI tissue-associated microbiota composition in 10 baboons using 16S rRNA amplicon sequencing (16S-Seq). We found the small intestinal (duodenum, jejunum and ileum) microbiota had significantly lower phylogenetic diversity (p < 1 × 10^−7^, two-tailed t-test, **Figure 1A**), lower Shannon indices (p < 0.0005, **Figure 1B**) and lower Chao1 indices (p < 1 x 10^−6^, **Figure 1C**) as compared to the microbiota in the large intestine (cecum, proximal colon and distal colon) [21, 26].

**Figure 1:**
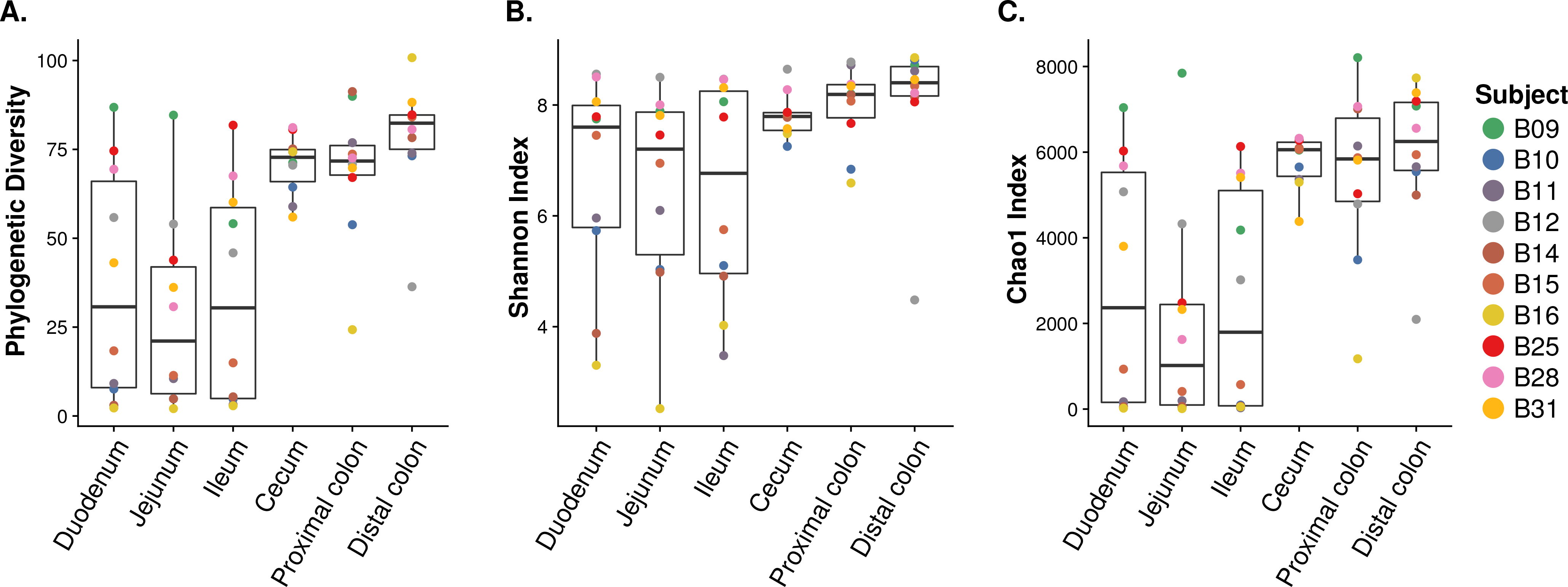
Microbiota alpha diversity along the GI tract. **a.**Phylogenetic diversity, **b.** Shannon Index, and **c.** Chao1 Index.

Microbial communities in the small and large intestines were predominantly comprised of members of the phyla Firmicutes, Bacteroidetes, and Proteobacteria, the relative abundances of which did not vary significantly between locations (Kruskal-Wallis test False Discovery Rate adjusted *P* ≥ 0.653 **Figure 2A**). However, overall community composition did vary between the small and large intestines (PERMANOVA p < 0.001; **Supplementary Figure 1**), and abundances of the less abundant phyla Spirochaetes, Tenericutes, and Lentisphaera were significantly greater in the large intestines (Kruskal-Wallis test FDR-adjusted *P* = 0.0275, 0.0137, and 0.0137, respectively; **Figure 2A**). Similarly, when classified to genus (**Supplementary Figure 2**), abundances of the genera *Flexispira, Sphaerochaeta, Anaerostipes,* and *Roseburia*, were greater in large intestines (Kruskal-Wallis test FDR-adjusted *P* = 0.0028, 0.0028, 0.0327 and 0.0837, respectively; **Supplementary Figure 3**).

**Figure 2:**
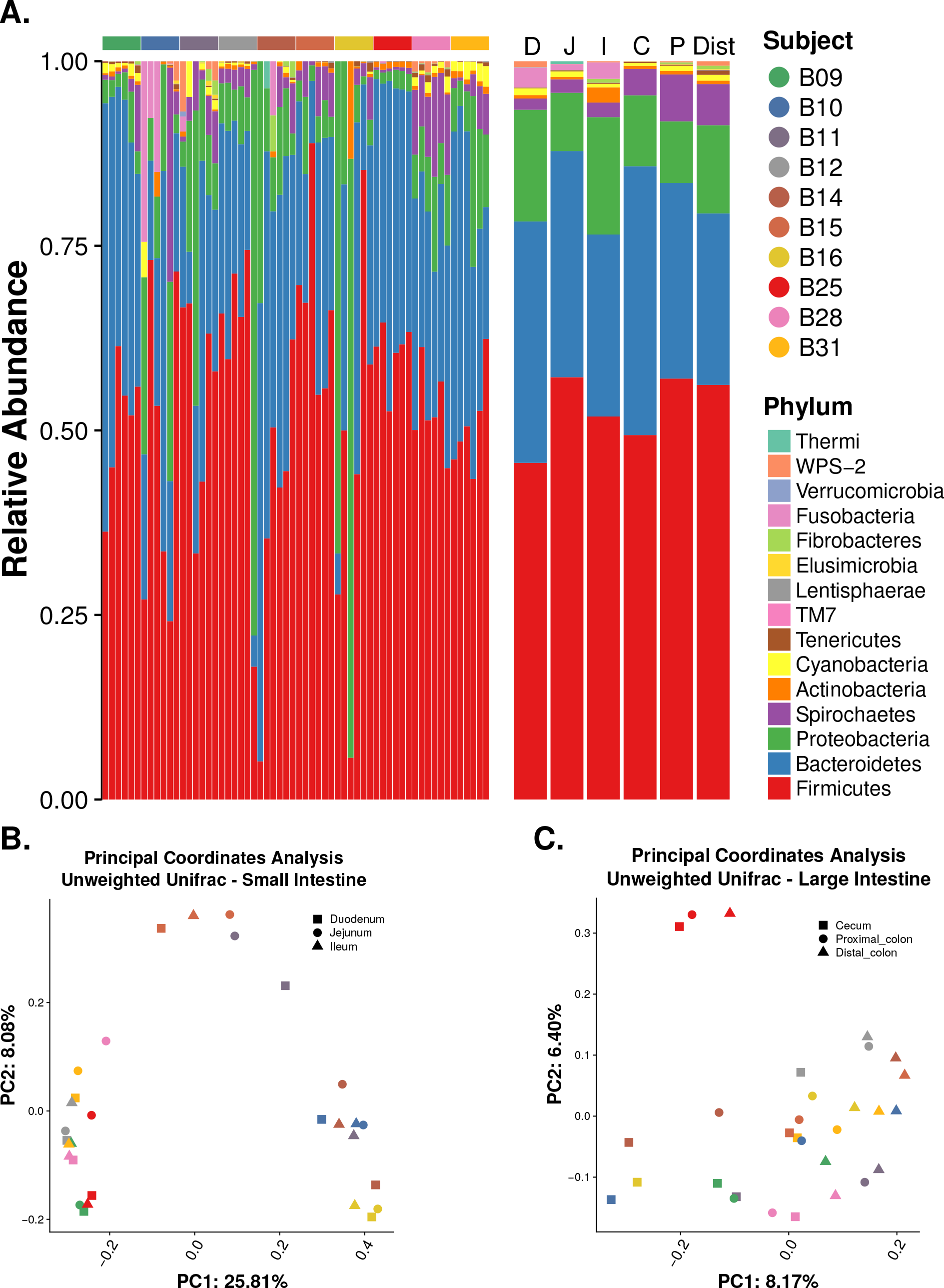
Microbiota along the non-human primate gastrointestinal tract. Stacked bar plot of bacterial phyla showing the relative abundance of **a.** each of the 6 tissue locations for each sample and the average for each of the 6 tissue locations. Unweighted UniFrac Principal Coordinates Analysis (PCoA) of **b.** small intestines and large intestines.

Among tissues in the small intestines, no significant differences were observed among specific sites (PERMANOVA P > 0.8; **Figure 2B**), but only one OTU classified to the family *Lachnospiraceae* had significantly greater relative abundance in the ileum, as determined by LEfSe (LDA = 2.68, *P* = 0.017; **Supplementary Table1 1**). Microbial communities among the small intestinal locations instead showed significant differences by individual subjects (PERMANOVA, *P* < 9.999e-05), but pairwise comparisons among individuals were not significant at Bonferroni-corrected α = 0.001. Conversely, communities among large intestinal tissues differed significantly by location (PERMANOVA, *P* < 0.001; **Figure 2C**), with significant differences noted between the cecum and distal colon (*P* < 0.001). Specifically, cecal communities showed greater relative abundances of OTUs classified within the *Prevotellaceae* (LDA = 3.06 – 4.36, *P* ≤ 0.042; **Supplementary Table 1**), while distal colon communities showed greater abundances of OTUs within the *Ruminococcaceae* (LDA = 3.02 – 3.87, *P* ≤ 0.020; **Supplementary Table 1**). Similar to the small intestine tissues, individual differences were also broadly observed in the large intestines (PERMANOVA, *P* < 0.05) but pairwise comparisons were not significant after correction for multiple comparisons.

### Metabolomic compositions along the baboon GI tract

Using the untargeted metabolomics method, we analyzed the tissue metabolome composition in tissue samples immediately adjacent to the tissues used for 16S rRNA gene sequencing. A total of 3,395 features (presented as retention time and mass to charge ratio) were identified in at least two-thirds of all the samples analyzed. After searching against the Human Metabolome Database (HMDB) and in-house libraries generated by the University of Minnesota Center for Mass Spectrometry and Proteomics, a total of 292 compounds were assigned identity. In our analysis, we focus on these compounds with assigned identities.

Due to broad differences in function and microbiome profiles between the small and large intestines, we sought to first identify differential metabolites between these tissue locations. We performed the Wilcoxon rank sum test and identified 140 metabolites with differential abundance between the small (87) and large (53) intestines (**Supplementary Table 2**). Among the 87 metabolites with significantly higher abundance in the small intestines, 24 are classified as amino acids, peptides, and analogs, such as aspartic acid, alanine, tyrosine, valine, leucine, and isoleucine. Long chain fatty acids such as linoleic acid and docosahexaenoic acid also have a higher presence in the small intestines. Additionally, Tauro-conjugated bile acids also have higher abundance in the small intestines [27, 28]. In the large intestine, there are more cholic acid and urobilin, in addition to other metabolites such as lysophosphatidylcholine and sphingomyelin.

To explore which metabolic pathways that the metabolites involve in, we performed pathway analysis using the differentially abundant metabolites between the small and large intestines using Ingenuity Pathway Analysis (IPA; **Supplementary Table 2**). We found that the differentially abundant metabolites are involved in the activation of bacterial growth related pathways (*P* = 3.36 x 10^−08^) in the small intestines. In the large intestines, the differentially abundant metabolites are involved in the activation of amino acids uptake pathways (*P* = 6.09x 10^−08^) and cancer-related pathways (*P* = 8.41Ex 10^−11^).

### Microbiota-Metabolome interactions

Next, we ask the question of whether there is any global microbiota-metabolome relatedness in the intestines. We performed the Procrustes analysis using the *vegan* package in R (**Figure 3**; **Supplementary Figure 4**). Globally, the Procrustes analysis shows significant relatedness (p = 0.039) between the microbiota and the metabolome. Interestingly, this relatedness is only driven by the distal colon (p = 0.0088; **Supplementary Figure 4**). We then analyzed the microbiota-metabolites relationships using Spearman’s ranked correlation on the metabolites with assigned identity. We first summarized the microbiota OTU data to the lowest level with a known bacterial assignment, then performed the correlation with the corresponding metabolites data. Globally, without regard to the tissue location, a total of 4,410 significant correlations (q < 0.1, False discovery rate adjusted p-value) were found between the microbiota and the known metabolites. Interestingly, when breaking the correlations down to the small and large intestines, the small intestine had 315 significant interactions (**Figure 4A**) while the large intestine had 139 (**Figure 4B**). Additionally, the correlation network in the small intestine is more interconnected as compared to the large intestine. Indeed, when we performed a similar analysis using the phyla level data, the large intestine had 78 significant correlations (q < 0.1) compared to 39 significant correlations (q < 0.1) for the small intestine. Interestingly, both networks appeared to have a microbiota-centric correlation, meaning most metabolites only correlated with relative abundances of a few taxa and such correlations tend to be in the same direction, however, one taxon correlated to many metabolites in different directions.

**Figure 3:**
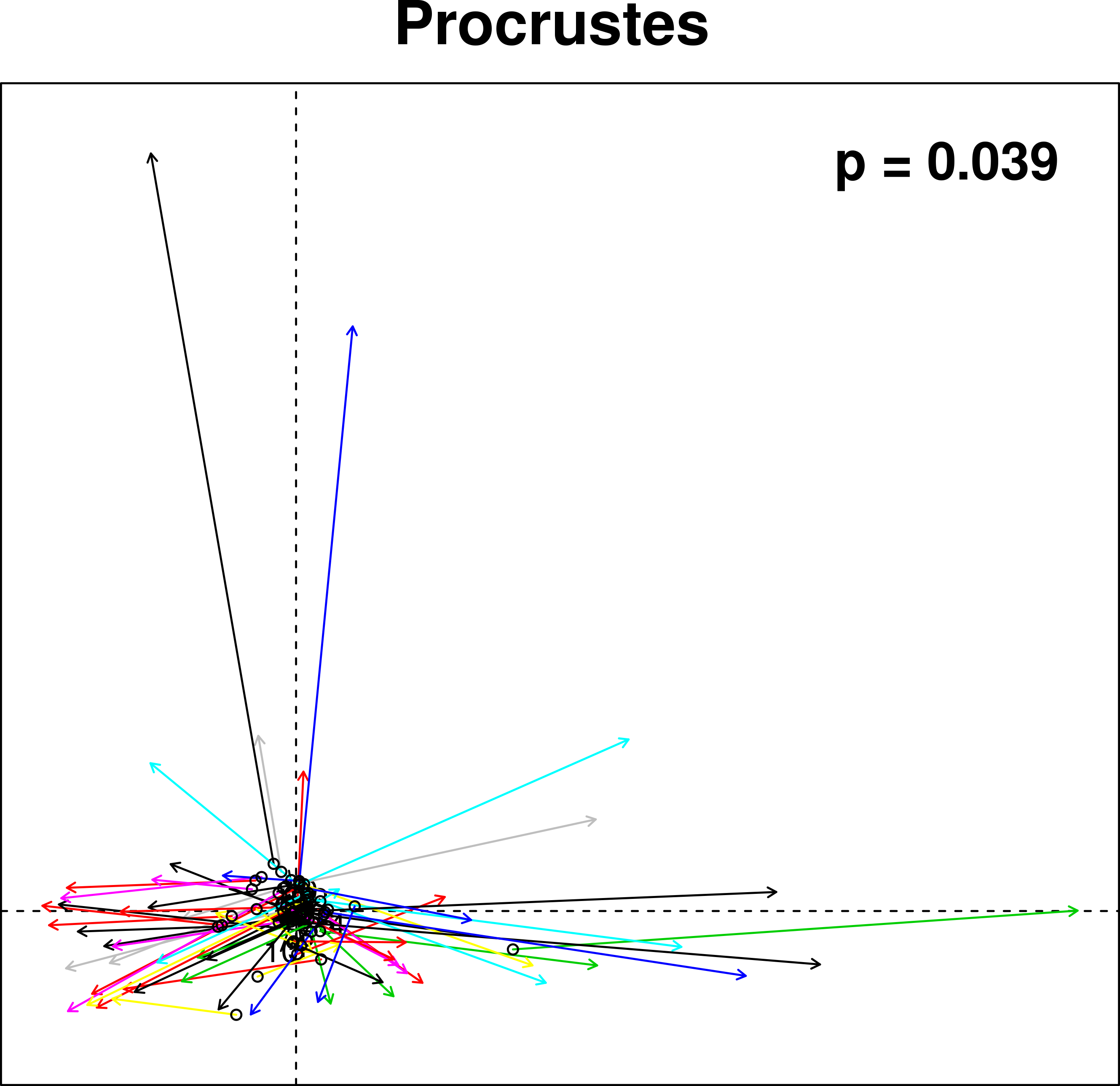
Procrustes analysis of the microbiota PCA against the metabolome Principal components analysis (PCA). Longer line length indicates less within sample similarities.

**Figure 4:**
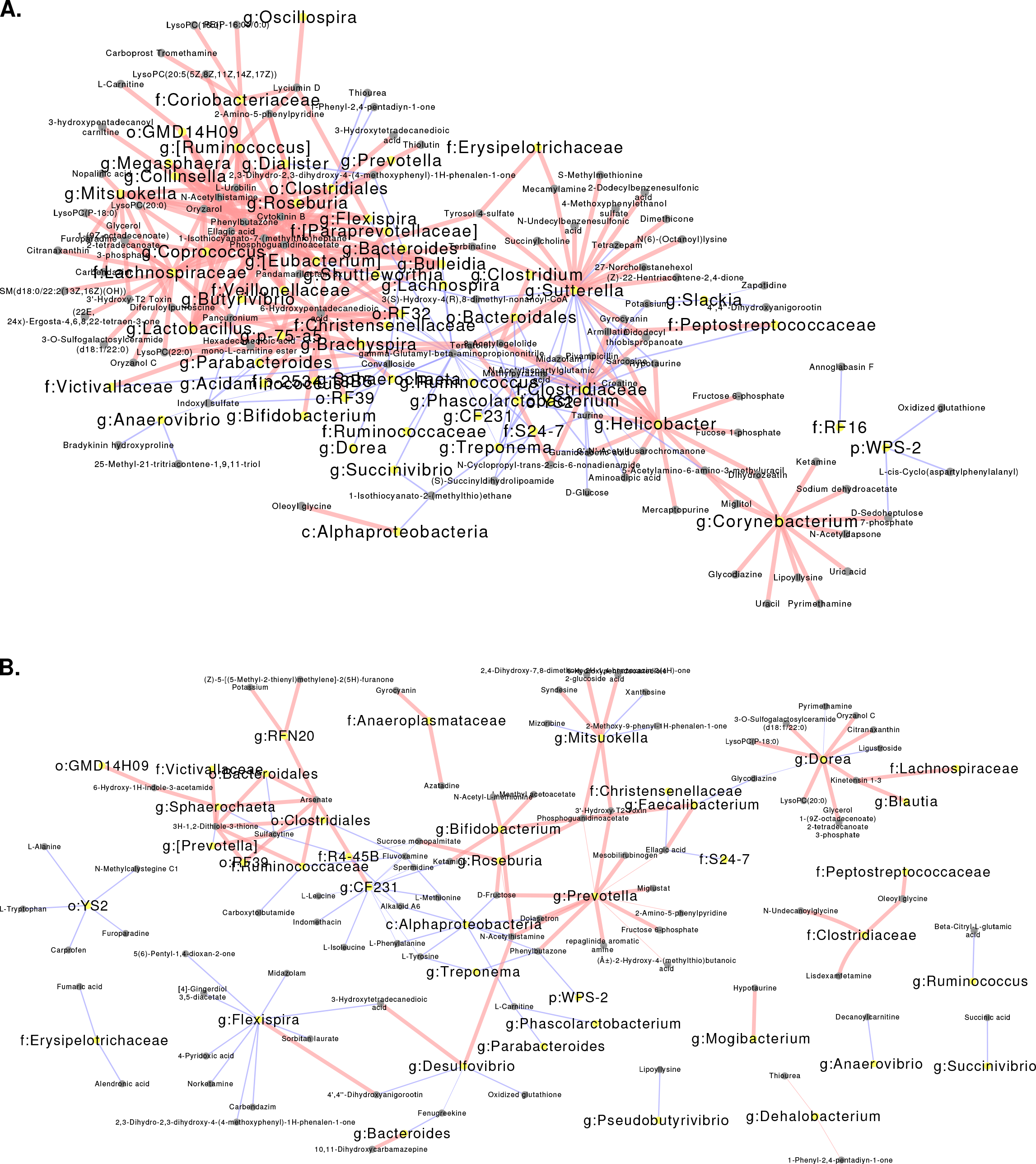
Microbiota-metabolome correlation network. Significant Spearman’s correlations of the **a.**small and **b.** large intestine, were visualized using Cytoscape. Red line indicates positive correlations and blue line indicate negative correlations. The thickness of the link indicates significance, where thicker link indicates more significant correlations.

The metabolites with the most bacterial correlations in the small intestines are a long-chain fatty acid, 6-hydroxypentadecanedioic acid, and the isothiocyanate, 1-isothiocyanato-7-(methylthio)heptane, with 19 and 16 significant correlations in the small intestines, respectively (**Figure 5A**). The sulfur-containing dithiolethione, 3H-1,2-dithiole-3-thione, had 7 significant correlations in the large intestine (**Figure 5B**). Interestingly, all 3 compounds are commonly found in brassicas vegetables, which were fed to the baboons as a part of the normal dietary enrichment that all animals received. All 19 bacterial taxa, including *Christensenellaceae*, *Bifidobacterium*, and *Lactobacillus*, correlated with 6-hydroxypentadecanedioic acid in the small intestine had positive correlations with this compound. These data warrant further investigation to elucidate the causal relationship between the mucosal microbiota and metabolites composition change.

**Figure 5:**
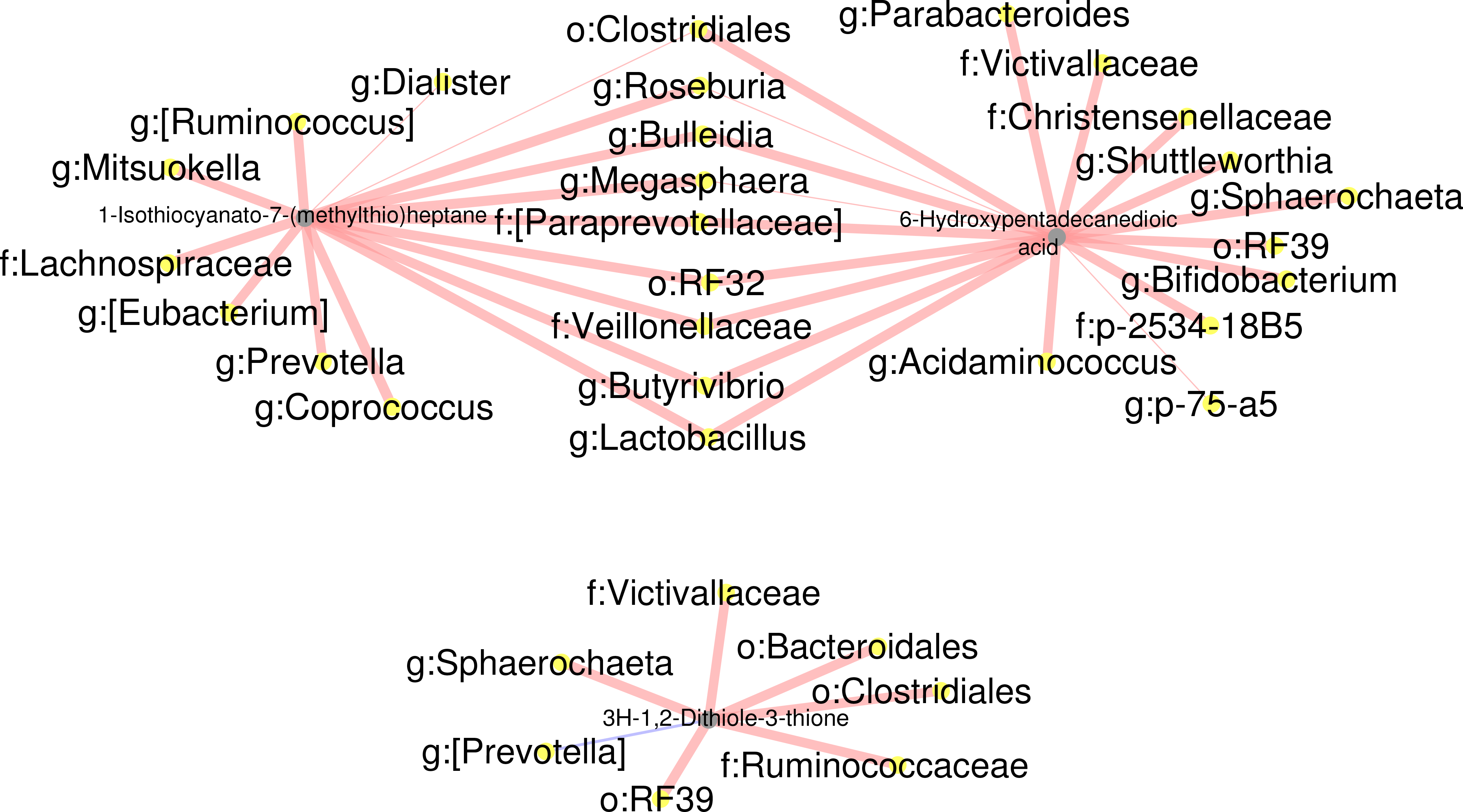
Microbiota-metabolites correlations. Significant Spearman’s correlations between the microbiota and **a.** 6-Hydroxypentadecanedioic acid, 1-Isothiocyanato-7-(methylthio)heptane, and **b.** 3H-1,2-Dithiole-3-thione were visualized using Cytoscape. Red line indicates positive correlations and blue line indicate negative correlations. The thickness of the link indicates significance, where thicker link indicates more significant correlations.

## Discussion

Currently, there is limited knowledge regarding the microbiota composition along different sections of the GI tract in either human or NHP samples [29, 30]. Studies in human subjects usually require prior bowel preparation, which has been shown to alter the microbiota [31]. In this study, we collected tissue samples from healthy, non-human primates without prior bowel preparation, thus providing an unaltered view of the healthy microbiota. Previous studies have analyzed the GI tract microbiota compositions in mouse, chicken, dog, cow, and horse [21–25]. However, due to the anatomical differences from humans, in addition to the dietary and genetic differences, these animals likely have different microbiota along the GI tract. To our knowledge, this study is the first report to comprehensively analyze the mucosa-associated microbiota-metabolite interactions along the GI tract in baboons.

Perhaps not surprisingly, the baboon intestinal microbiota composition is more similar to that observed in human GI tissue-associated microbiota and differs from that observed in mouse fecal samples (**Supplementary Figure 5**) [5]. Contrary to a previous human study which examined the microbiota composition using small and large intestinal biopsy samples, we found limited microbiota differences along the intestinal tract at the phyla level (**Supplementary Figure 5**) [26]. Additionally, we found lower alpha diversity in the small intestines, while Stearn et al. did not. One plausible explanation is the previous study collected biopsy samples after the patients went through bowel preparation, and this may affect the microbiota composition. Indeed, the fecal samples collected prior to the bowel preparation had very different microbiota composition compared to the colon tissues in the previous study [26].

Similar to previous reports in humans, we found variations to the microbiota composition between different NHP subjects. Previous studies suggest that this variation between individuals can be attributed to factors such as genetics, dietary preferences and other factors [1, 32, 33]. We found that both small and large intestinal tissues have significant interindividual variations, although the statistical significance in the small intestinal tissues is more pronounced compared to the large intestines (PERMANOVA, *P* < 9.999e-05 vs. *P* < 0.05). This together with the differences we observed in alpha diversity along the GI tract sections is likely due to the microbial concentration gradient along the GI tract, where the small intestine harbors lower concentrations of bacteria due to the high pH environment [34]. It is also not surprising that the distal colon and cecum harbor more distinct bacterial taxa compared to other locations. Previous studies have shown that both distal colon and the cecum are where most bacterial fermentation takes place, although we found no discernable differences in the predicted composition of the microbiota [35].

Interestingly, we found that the host-microbiota metabolic interactions in the intestines are microbiota-centric, with most metabolites correlated with only a few taxa, and in the same direction. It is important to note that this analysis could not identify the directionality of such interactions; however, these data may suggest that the microbiota exerts a greater influence on metabolic interactions than is regulated by the host. Interestingly, only 4 microbiota-metabolome correlation pairs were found in common between the small and large intestine. One explanation is due to the higher level of functional redundancy present in the large intestine, as the result of more species present. Another possible explanation is that environmental variables including pH and oxygen availability, as well as the nutrient compositions are different between the small and large intestines [34, 36]. These factors together may have additional selective effects on the microbiota and the tissue metabolome.

Based on the microbiota-metabolites correlation network, we unexpectedly found that 6-hydroxypentadecanedioic acid, 1-isothiocyanato-7-(methylthio)heptane, and 3H-1,2-dithiole-3-thione, compounds commonly found in brassica vegetables, were correlated with greater relative abundances of several bacteria previously shown to produce health benefits in humans. They include *Christensenellaceae*, *Bifidobacterium* and *Lactobacillus [37, 38]*. This limited information suggests that 6-hydroxypentadecanedioic acid may have potential prebiotic effect. Furthermore, 3H-1,2-dithiole-3-thione has been previously shown as a potent antioxidant and potential chemo-preventative agent [39]. This supports the potential prebiotic effect of this compound. Moreover, these compounds show location specific correlations with microbiota, which may suggest a potential strategy to target beneficial bacteria in different intestinal locations. One explanation to the location specific correlations is the differences of absorptions of these compounds at different locations of the intestine which can lead to different metabolite concentrations in the intestinal lumen. However, since the current study did not include controlled feeding, we are unable to ascertain the exact role of these compounds in modulating the microbiota. Considering the potential health benefits associated eating brassicas vegetables, this finding warrants additional investigations.

In this study, we performed untargeted metabolomics on the intestinal tissues. Although we were able to identify over 3,395 compounds, we were able to assign identities to 292 compounds. This lack of positive identification is mainly due to under-representation of metabolites in available databases. It is conceivable that future database expansion will allow us to extract additional information from the data. Additionally, the current research is unable to distinguish between the metabolic contribution from the host and the microbiota. Thus, whether this metabolic microenvironment differences between the small and large intestines is solely contributed by the microbiota is yet to be determined. The previous study comparing the tissue level metabolome between conventional and germ-free mice showed that the microbiota contributes to various metabolomic differences along the GI tract. However, whether this difference is due to changes to the microbial metabolism or host metabolism is unclear [28]. Future studies should aim to separate metabolites originated from the host, microbiota or food source.

## Conclusions

In the present study, we report the microbiota and metabolomic profiles of the small and large intestinal sections of healthy non-human primate (NHPs). Our study provided a global view of the intestinal microbiota landscape of healthy NHPs and revealed an intricate global relationship between the microbiota and the metabolites along the GI tract. Our analysis disseminated the intestinal location specific microbiota and metabolites correlations and provided evidence that warrants future functional validation to establish the directionality of such interactions.

## Methods

All protocols and procedures were approved by the University of Minnesota Institutional Animal Care and Use Committee, conducted in compliance with the Animal Welfare Act, and animals were housed and cared for according to the standards detailed in the Guide for the Care and Use of Laboratory Animals. The cohort included 10 adult purpose-bred female olive baboons *(Papio anubis)* modeling ACL injury and subsequent repair using regenerative medicine techniques. Animals were between 6.5 and 15.6 years (median, 9.3 years) in age and weighed between 14.4 and 24.9 kg (median, 20.1 kg). They were pair housed or housed in protected contact with compatible conspecifics. Baboons had free access to water and were fed identical diets that included biscuits (Harlan Primate Diet 2055C, Harlan Teklad) based on body weight and daily enrichment with fresh fruits, vegetables grains, beans, nuts, and a multivitamin preparation. Semiannual veterinary physical examinations were performed in all animals. Animals participated in an environmental enrichment program designed to encourage sensory engagement; enhance foraging behavior and novelty seeking; promote mental stimulation; increase exploration, play, and activity levels; and strengthen social behaviors, together providing opportunities for animals to increase the time budget spent on species-typical behaviors. Baboons were trained to cooperate in medical procedures including hand-feeding and drinking, shifting into transport cages for sedation and targeting or presentation for examination. Animals were euthanized via barbiturate overdose (Beuthanasia-D ≥86 mg/kg IV) and tissue procurement performed post mortem. No oral medications were used for at least 6 months prior to tissue collection. Approximately 1cm × 1cm tissue sections that included duodenum, jejunum, ileum, cecum, proximal colon, and distal colon were collected using clean technique and snap-frozen in liquid nitrogen and then stored at −80°C.

### 16S rRNA gene sequencing and sequence analysis

Total DNA was extracted from approximately 250 mg of tissue using the DNeasy PowerSoil Kit (Cat: 12888; Qiagen, Valencia, CA) following the standard protocol. Sequencing libraries were created by the Mayo Clinic Genome Analysis Core (Rochester, MN). Briefly, the V3-V5 region of the 16S rRNA gene was amplified with multiplexing barcodes using PCR (V3-341F: TCGTCGGCAGCGTCAGATGTGTATAAGAGACAGCCTACGGGAGGCAGCAG; V5-926R: GTCTCGTGGGCTCGGAGATGTGTATAAGAGACAGCCGTCAATTCMTTTRAGT). The libraries were then pooled and size-selected between 700 and 730 bp using a LabChip XT (PerkinElmer, Waltham, MA). Sequencing was performed on a single lane of a MiSeq sequencer (Illumina) using paired-end mode at a read length of 150 nucleotides. On average, 64,937 quality reads (between 9,901 and 118,288) were generated per library. The sequencing results were analyzed using the gopher-pipelines (https://bitbucket.org/jgarbe/gopher-pipelines) developed and maintained by the University of Minnesota Informatics Institute. Briefly, the adapters and low-quality reads were first trimmed using Trimmomatic v0.33. Then, the forward and reverse read pairs were merged using PandaSeq v2.8 [40]. OTUs were then picked using QIIME v1.9.1 “pick_open_reference_otus.py” script against Greengenes 16S database (May 2013 release), allowing 97% similarity [41, 42]. The unfiltered OTU table is available in **Supplementary Table 3**. The beta-diversity between tissue locations were analyzed by performing Principal Coordinate Analysis (PCoA) using both weighted and unweighted UniFrac distance metrics [43].

### Metabolites extraction

Metabolites were extracted from the immediate adjacent tissue that was used to generate 16S rRNA sequencing. There was an insufficient amount of duodenum tissue from animal B09 to perform untargeted metabolomics, and thus this was not analyzed. Approximately 15 mg of tissue was used to extract metabolites. The tissues were first ground into fine powder using a CryoGrinder (OPS Diagnostics) on dry ice. The tissues were then suspended in 20 µl of 80% methanol per 1 mg tissue weight. The mixture was then homogenized using a probe sonicator at 10% amplitude for 15 s, with 1-minute rest on ice between every 5 s of sonication. The sonicated samples were then centrifuged at 14,000 × *g* for 10 min at 4 °C. The supernatant from the centrifugation contained the metabolites and was saved in −80 °C. The tissue pellets were then further processed for additional metabolites extraction. They were first suspended in 10 µl of 80% methanol per 1 mg of original tissue weight and sent through high-pressure cycling on a Barocycler NEP2320 (Pressure Biosciences). The high-pressure cycling protocol include 60 cycles of 20 s of 35,000 psi pressure, followed by 10 s of 0 psi for at 4 °C. After pressure cycling, the samples were again centrifuged at 14,000 × *g* for 10 min at 4 °C and the supernatant was pooled with the previously extracted metabolites. Finally, the metabolites were dried under a nitrogen stream.

### Untargeted metabolomics

The dried metabolites were first suspended in 15 µl of 0.1% formic acid per 1 mg of the original tissue weight. The suspensions were then separated for analysis using a C18 reverse-phase column and hydrophilic interaction liquid chromatography (HILIC) column. The reverse-phase analysis results in separation of larger non-polar molecules such as steroid-like compounds, certain amino acids, phospholipids and other lipids, while the HILIC analysis separates hydrophilic compounds such as amino acids and members of the citric acid cycle and glycolysis pathways. The samples were analyzed using reverse-phase positive mode (non-polar interaction) separation and HILIC analysis (polar interaction) separation before analyzed by Q Exactive LC-MS/MS quadrupole Orbitrap (Thermo Scientific). The reverse-phase analysis was performed in positive mode ionization with an additional proton (+1.0073) added. For HILIC analysis, the negative ionization mode was used with one additional proton (−1.0073) removed. Since salts are present, compounds may occasionally form as a sodium salt (neutral mass plus 21.9944) for positive or a chloride salt (neutral mass plus 34.9688) for negative mode. Samples were loaded and analyzed in random orders and quality control samples were analyzed in regular intervals to eliminate extraneous signals. The untargeted metabolomics were performed by the University of Minnesota Center for Mass Spectrometry and Proteomics.

### Metabolomics data analysis

The data was processed using Progenesis QI software (Thermo). The software first aligns all the features obtained in all the runs and then assigns intensity measures for features found in all the runs. The raw data were further processed by filtering for fidelity of individual feature detection using the quality control samples. Only features with a CV less than 10% among all quality control samples were accepted. Features showing high intensity in background samples relative to the quality control samples and features not present in at least 67% of all samples were removed from analysis as per U.S. Food and Drug Administration recommendation. Each feature is uniquely identified with the mass-to-charge ratio (m/z) and the elution time from the column. Features were then assigned to metabolites identified by searching the Human Metabolome DataBase (HMDB) and using databases developed by the University of Minnesota **(Supplementary Table 4)**. Pathway analysis was performed using Ingenuity Pathway Analysis (IPA).

### Microbiome-Metabolome correlation analysis

The Spearman’s ranked correlation test with false discovery rate (FDR) adjustment were used to test the microbiome-metabolome correlation [44]. The microbiome OTU data and metabolomics data were first combined and filtered to remove low abundance OTUs and metabolites (appearing in less than 50% of samples). The Spearman’s ranked correlation were calculated using *cor.test* function in R v3.4.4. The p-values were then adjusted using *p.adjust* function before filtering for significant correlations. The Spearman’s rank correlation is a nonparametric test that can be used to reveal a subtle relationship between microbiota and metabolites [44].

## Declarations

### Ethics approval and consent to participate

All protocols and procedures were approved by the University of Minnesota Institutional Animal Care and Use Committee, conducted in compliance with the Animal Welfare Act, and animals were housed and cared for according to the standards detailed in the Guide for the Care and Use of Laboratory Animals.

### Consent for publication

Not applicable

### Availability of data and material

All data generated or analyzed during this study are included in this published article and its supplementary information files. The 16S-Seq data are available on Sequence Read Archive (SRA) under the assession number SRAxxxxxx.

### Authors’ contributions

C.Y. and S.S. developed the concept, C.Y. carried out data analysis with assistance from C.S. and S.S. C.Y., M.G., C.S. and S.S. wrote the manuscript.

## Acknowledgments

The authors thank the members of the Subramanian lab and the University of Minnesota Center for Mass Spectrometry and Proteomics (CMSP) for helpful discussions. We thank the Preclinical Research team and Research Animal Resources veterinarians for providing outstanding animal care and for providing tissue via the NHP tissue sharing program in an effort to reduce the overall number of NHPs used in research. We thank Dr. John Garbe of University of Minnesota Research Informatics Service for developing the gopher-pipelines. We also thank the Medical Genome Facility Genome Analysis Core at Mayo Clinic (Rochester, MN) for performing library prep and sequencing and CMSP for performing the untargeted metabolomics. This work was carried out, in part, using computing resources at the Minnesota Supercomputing Institute.

## Supplementary Data

**Supplementary Table 1:** Linear discriminant analysis (LDA) effect size (LEfSe) analysis of small and large intestinal microbiota.

**Supplementary Table 2:** Differentially abundant metabolites between small and large intestines and Ingenuity pathway analysis.

**Supplementary Table 3:** Unfiltered OTU table used for analysis.

**Supplementary Table 4:** Unfiltered untargeted metabolomics data used for analysis.

**Supplementary Figure 1:** Principal Coordinates Analysis (PCoA). **a.** weighted UniFrac PCoA and **b.** unweighted UniFrac PCoA showing all samples. Weighted UniFrac PCoA of **c.** small intestines and **d.** large intestines.

**Supplementary Figure 2:** Stacked bar plot of bacterial genera showing the relative abundance of the 6 tissue locations.

**Supplementary Figure 3:** Tissue specific procrustes analysis. **a.** Jejunum, **b.** Duodenum, **c.** Ileum, **d.** Cecum, **e.** Proximal Colon, and **f.** Distal Colon. PCA of the **g.** tissue metabolome and **h.** microbiota.

**Supplementary Figure 4:** Ingenuity Pathway Analysis of differentially abundant metabolites. Pathways related to **a.** Growth of bacteria is activated in the small intestines. Pathways related to **b.** Uptake of amino acids and **c.** solid tumor is activated in the large intestines. The blue line indicates activation, the orange line indicates inhibition, yellow line indicates conflicting evidence, the grey line indicates an association.

**Supplementary Figure 5:** Stacked bar plot of bacterial phyla showing the relative abundance comparing the Human, Baboon and Mouse microbiota samples.

