## Supplementary figures and images for "Mucosal microbiota and metabolome along the intestinal tracts reveals location specific relationship"

### Supplementary Figure 1

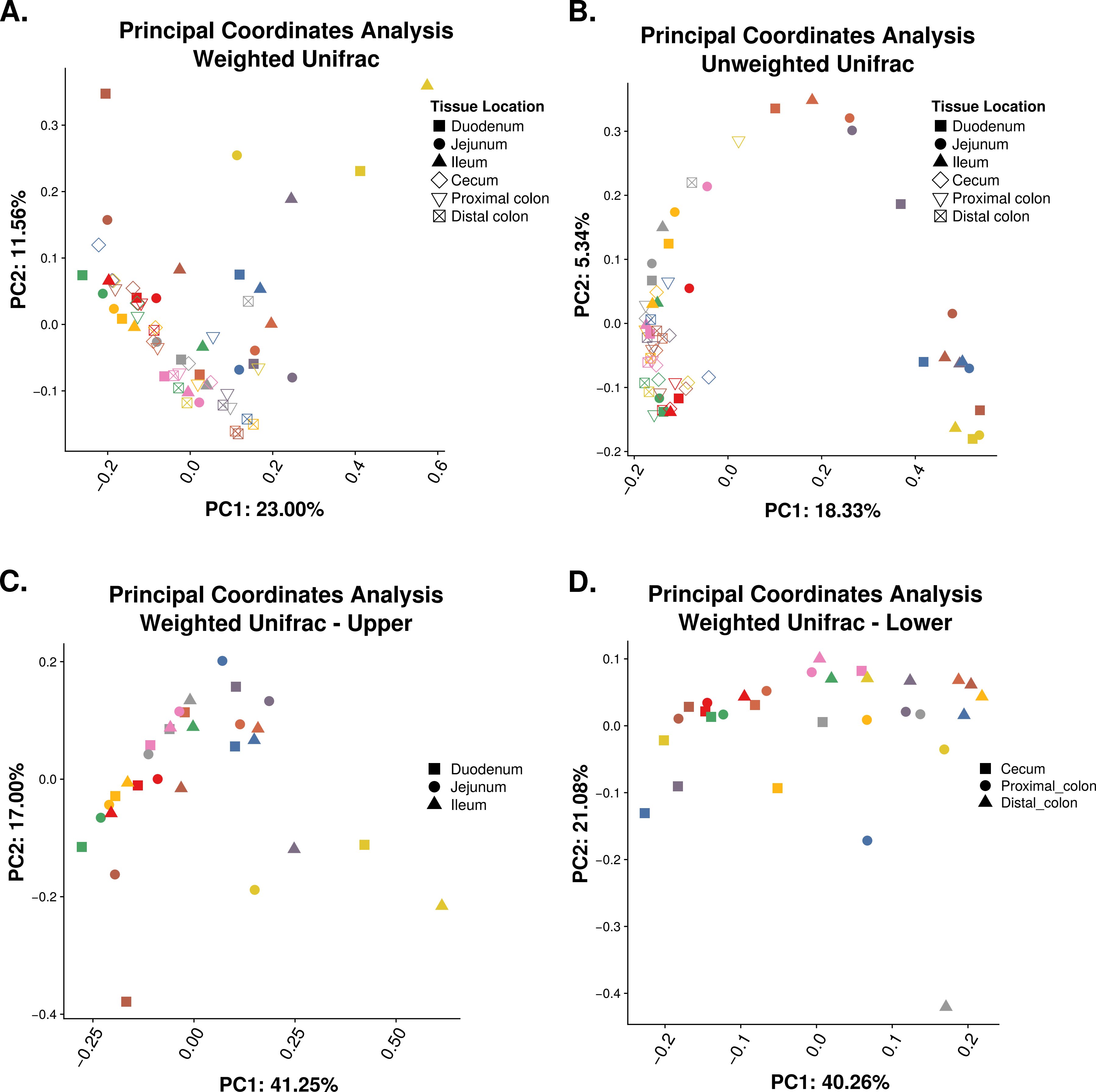

### Supplementary Figure 2

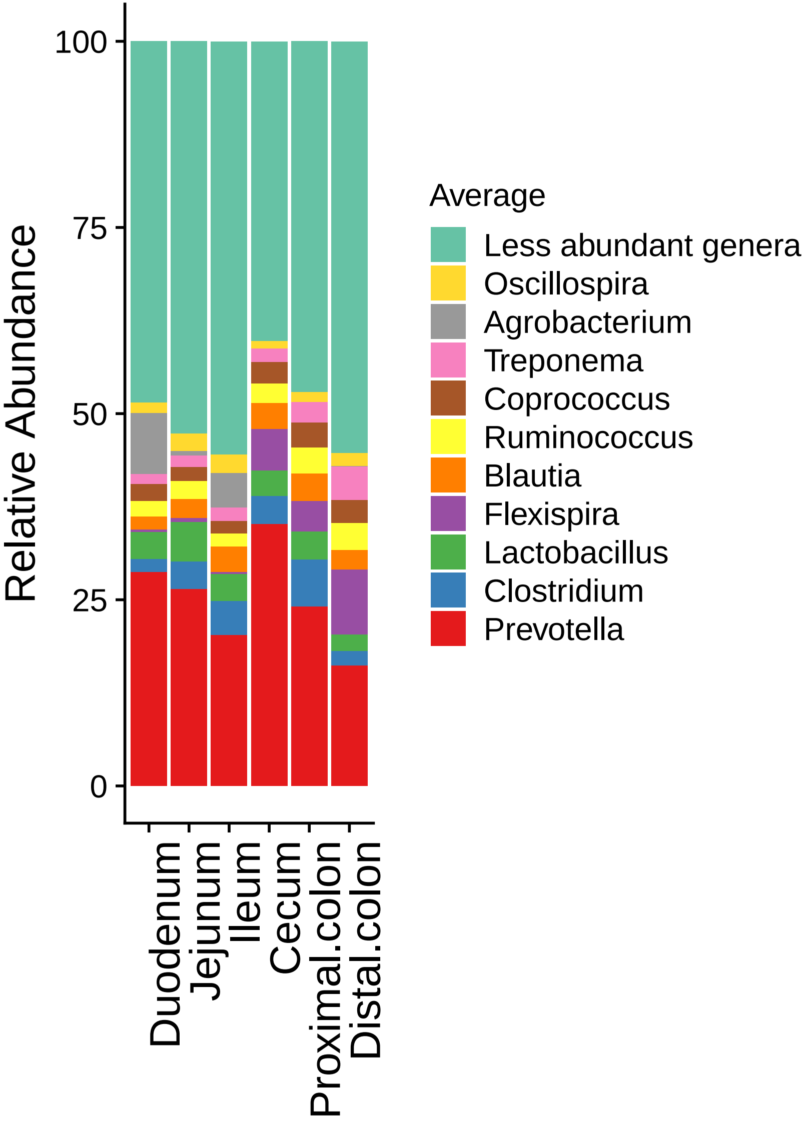

### Supplementary Figure 3

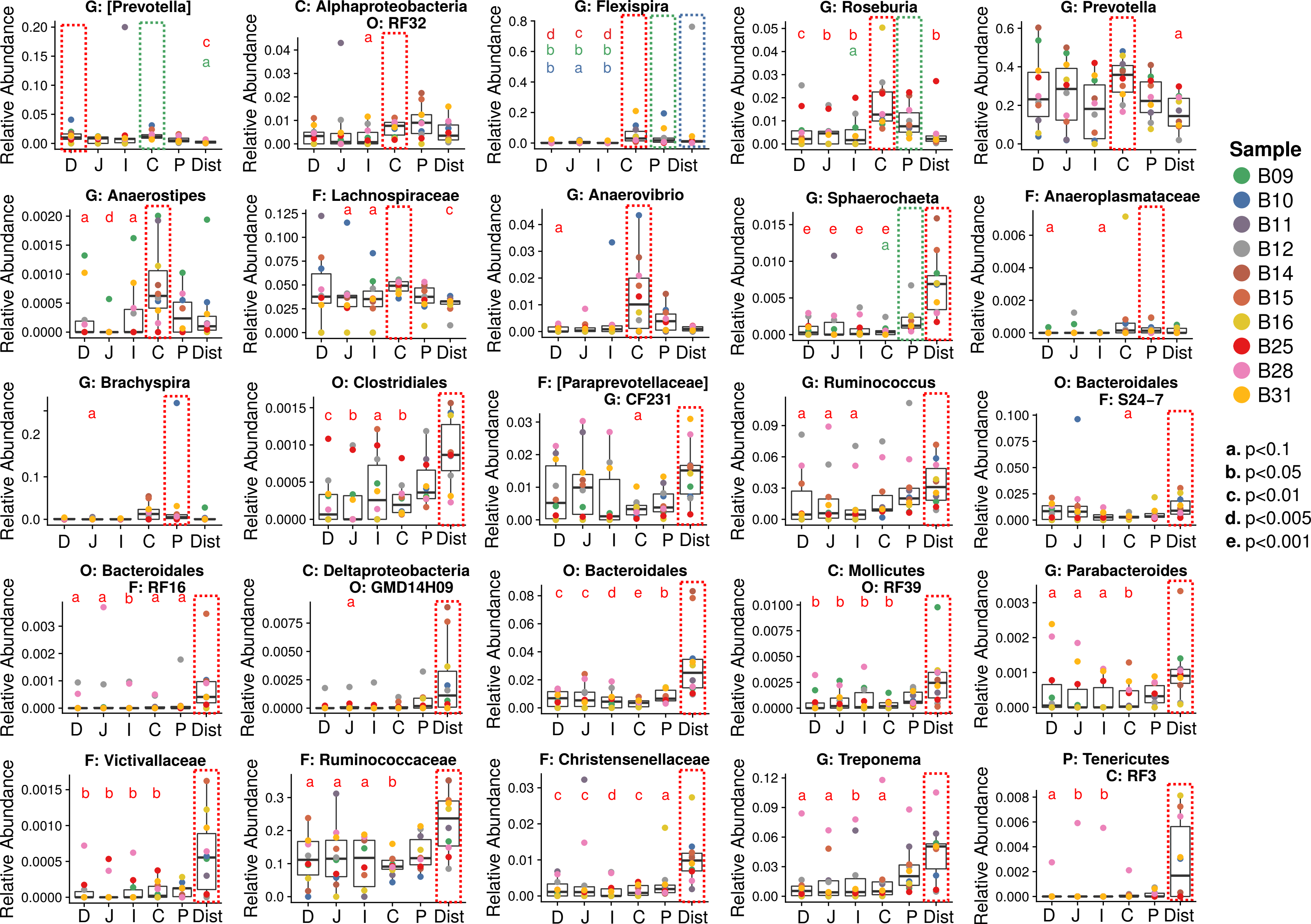

### Supplementary Figure 4

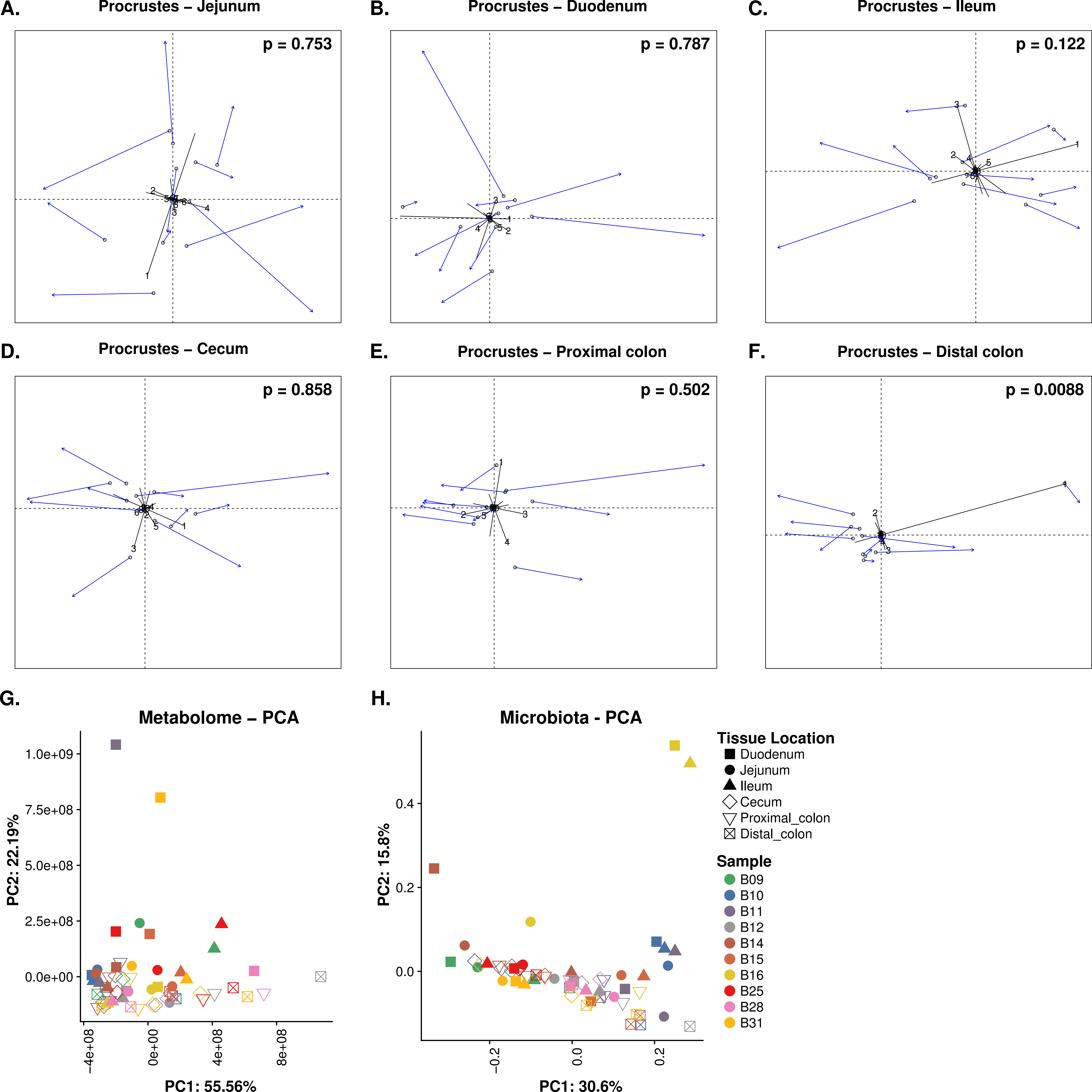

### Supplementary Figure 5

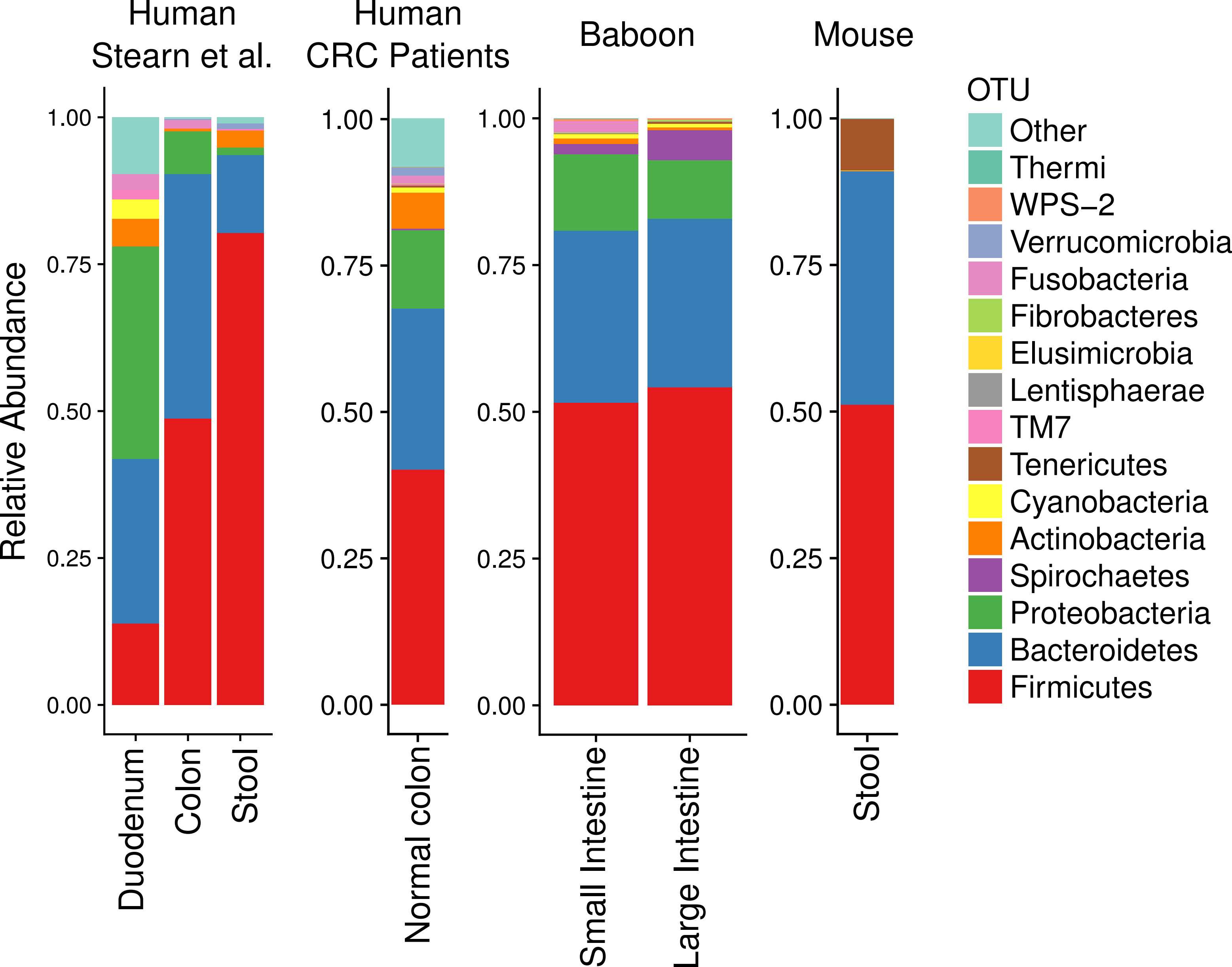
